# Putative long-range mossy fiber sprouting and regional hypermetabolic capacity in the hippocampus of patients with mesial temporal lobe epilepsy

**DOI:** 10.1101/2025.07.17.665452

**Authors:** Tian Tu, Lily Wan, Qi-Lei Zhang, Chen Yang, Hong-Shu Zhou, Zhong-Ping Sun, Hong-Yu Long, Bei-Sha Tang, Aihua Pan, Ewen Tu, Jian Wang, Zhi-Quan Yang, Zhen-Yan Li, Xiao-Xin Yan

**Affiliations:** Department of Neurology, Xiangya Hospital, Central South University, Changsha, Hunan 410008, China; Department of Anatomy and Neurobiology, Xiangya School of Basic Medical Sciences, Central South University, Changsha, Hunan 410013, China; Department of Neurosurgery, Xiangya Hospital, Central South University, Changsha, Hunan 410008, China; Department of Neurology, The Second People’s Hospital of Hunan Province, Changsha, Hunan 410007, China; The Reproductive and Stem Cell Engineering Institute, Xiangya School of Basic Medical Sciences, Central South University, Changsha, Hunan 410013, China

**Author notes:** Correspondence to: Xiao-Xin Yan,; Zhen-Yan Li.

**Keywords:** Aberrant neuroplasticity, Actin isoforms, Epilepsy, Hippocampal sclerosis, Neuronal death, Tri-synaptic circuit

## Abstract

Mesial temporal lobe epilepsy (MTLE) is pathologically characterized by neuronal loss in the dentate hilus, CA3 and CA1 regions, and mossy fiber (MF) sprouting into the inner molecular layer (iML). The latter forms aberrant excitatory circuities that are considered to facilitate recurrent seizures, with the subiculum also being related to epileptogenic activation. We recently identified a distinct expression of α-smooth muscle actin (αSMA) at the MF terminals in human hippocampus. This prompted us to explore MF sprouting in resected hippocampi (n=20) from patients with MTLE relative to postmortem control (n=20) using αSMA along with reference markers for pathological cross-validation. Compared to control, neuronal loss assessed with neuron-specific nuclear antigen and sortilin immunolabeling reached CA1 in all resected hippocampi. αSMA, zinc transporter 3 and β-secretase 1 immunolabeling in the iML tended to be increased. The MF-related markers also revealed a preserved fibrous band extending across CA1 to subiculum. Cytochrome c oxidase immunolabeling also increased in iML and subiculum in the MTLE group. Taking together, the current findings point to the existence of long-range MF sprouting and a regional hypermetabolic compacity in the hippocampal formation of patients with drug resistant MTLE.

## Introduction

Mesial temporal lobe epilepsy (MTLE) is a devastating neurological disease, especially affecting children and young adults who are expected to fulfill immense life and career potentials^1^. MTLE can be progressive, with hippocampal sclerosis and relapsing seizures worsening to a certain point with symptoms becoming pharmacologically uncontrollable^2^. Neuronal loss, gliosis, granule cell dispersion and mossy fiber (MF) sprouting are the pathological characteristics of MTLE^3^. According to the pathological grading system of the International League Against Epilepsy (ILAE)^4^, neuronal loss and gliosis occur earlier and severely in the hilar region of the dentate gyrus (DG), CA3 and CA1, but can extend into CA2 and even the subicular subregions as sclerosis progresses. Granule cell dispersion involves increased cellular tiers and dislocation, reduced cell density and laminar disorganization. MF sprouting into the inner molecular layer (iML) has been documented with Timm stain and immunolabeling of various presynaptic markers such as zinc transporter 3 (ZnT3), dynorphin, vesicular glutamate transporters, synaptic vesicle proteins and β-secretase 1 (BACE1)^5–13^.

The sprouting MF terminals can form aberrant asymmetric synapses with the dentate granule cells, surviving hilar mossy cells and interneurons^14–18^. The reorganized MF terminals in the iML can develop the so-called perforated synapses, which are considered highly efficacious in synaptic transmission^19,20^. MF sprouting has been suggested to facilitate recurrent epileptiform activity in the DG^21,22^, while this has remained an issue of debate as blocking MF sprouting by rapamycin does not reduce seizure frequency in pilocarpine-treated mice^23^ and there also exists increased MF innervation to interneurons in the DG^24,25^. Studies in rodent models show that both neonatal and adult-born granule cells contribute to the formation of aberrant MF connections^26–29^ and that optogenetic activation of sprouted mossy fibers can detonate aberrant dentate network activity^30^.

The subiculum is relatively preserved during hippocampal sclerosis and has been considered to play a critical role in synchronization and propagation of epileptic activities into extrahippocampal brain regions^31^. Animal model and *ex vivo* human hippocampus studies have gained much electrophysiological and neuroanatomical evidence supporting this notion. The underlying mechanism includes enhanced excitatory and reduced inhibitory neuronal activity and altered glial modulation in this region^32–39^. The subiculum is therefore considered an area for potential pharmacological and surgical (such as deep brain stimulation) intervention for MTLE^40^.

We recently identified a novel α-smooth muscle actin (αSMA) expression in human hippocampal MF, which is highly selective relative to presynaptic markers that can display these giant synaptic structures^41^. The present study was set to explore MF pathology in resected hippocampi from patients with MTLE along with postmortem human brain samples as control using αSMA and a set of other markers for MF sprouting and neuronal loss^6,41^. To understand a potential metabolic basis for hippocampal hyperexcitability, we concurrently assessed the expression of cytochrome c oxidase (COX) in the resected samples relative to control by immunohistochemistry.

## Materials and methods

### Resected hippocampi and postmortem control samples

Human brain materials were used in the current study with approval (2020KT-37, 4/10/2020; #2023-KT084, 6/21/2023) by the Ethics Committee of Central South University Xiangya School of Medicine in compliance with the Code of Ethics of the World Medical Association (Declaration of Helsinki). All experimental procedures and methods were performed in accordance with the relevant guidelines and regulations set forth by the central south university. A total of 20 resected hippocampi were obtained from the Department of Neurosurgery of Xiangya Hospital following operation on patients with drug-resistant epilepsy and neuroimaging diagnosis of hippocampal sclerosis (Table 1). The samples were obtained between December 2020 and July 2022, with all patients remaining seizure-free to date. The surgical procedure and pathological examination of resected brain tissues were carried out with written consent from the patients and family members. The resected brain samples consisted of mostly the sclerotic hippocampus except for one case also including a part of the basal temporal cortex. The tissues available for this study were mostly a coronal slice at the mid-hippocampal level approximately in 0.5 cm thickness. The samples were placed on ice following resection, brought to a nearby pathological laboratory and dissected for use for this and other (ongoing) studies within approximately 30 minutes. The tissue slices used for the present study were fixed by immersion in formalin for 1-2 weeks.

Postmortem human brains were banked through a willed body donation program^42^. Each brain was bisected after removal from the skull, with a half cut into 1 cm-thick coronal slices, fresh-frozen then stored at -80 °C, and the other hemibrain immersed in formalin for 2-4 weeks and subsequently cut coronally into 1 cm-thick slices. Fixed brain blocks were prepared into cryostat (35 µm-thick) and paraffin (3 µm-thick) sections, followed by neuropathologically evaluation for the presence of Alzheimer’s and Parkinson disease pathologies according to the Standard Brain Banking Protocol proposed by China Brain Bank Consortium^43^. The postmortem brains were selected as control for the resected hippocampal group in an age/sex matchable manner, all of which lacked the above and other neuropathological lesions (Table 1).

### Tissue preparation, histological and immunohistochemical stainings

In general, the surgically removed hippocampi from patients and temporal lobe slices from postmortem human brains were cryoprotected following formalin fixation in 30% sucrose until tissue sank, embedded in optimal cutting temperature (OCT) compound, cut in a cryostat into 35μm-thick sections, which were stored in a cryoprotectant (30% sucrose, 30% ethylene glycol and 1% polyvinylpyrrolidone-40 (PVP-40 in 0.1 M phosphate buffer, pH 7.3) at -40 °C until use.

Cryostat sections were immunohistochemically stained in a batch-processing manner in 6-well tissue culture plate by including sections from 3-5 resected hippocampi and 3-5 postmortem brains in each experiment using the avidin-biotin complex (ABC) method (see Table 2 for antibody information). Selected sections were treated first in 0.1 M phosphate-buffer saline (PBS, pH 7.3) with 5% normal horse serum, 5% H_2_O_2_ and 0.1% Triton-100 for 1 hour (hr) at room temperature to block nonspecific reactivity, then reacted with a primary antibody at 4 °C overnight. On the second day, the sections were incubated in PBS containing biotinylated pan-specific secondary antibody for 1 hr and further reacted with the ABC reagent (1:200) for another hr at room temperature. Immunoreaction product was developed in PBS containing 0.05% diaminobenzidine (DAB) and 0.3% H_2_O_2_. A few sections were included in each experiment through all steps but the primary antibody incubation, which were used for defining background labeling. All the stained sections were mounted on glass microslides, allowed to air-dry, and coverslippered with a permanent mounting medium following dehydration and clearance in xylene.

### Image acquisition and densitometric analyses

The sections were scan-imaged with the 20× objective on a Motic-Olympus microscope using the same imaging setting. Images were examined on the interface of the Motic Digital Slide Assistant System Lite 2.0 (Motic Asia, Hong Kong, China), with the areas of interest at desired magnifications extracted for figure preparation. To avoid confusion with different uses of the term “CA4”, the area occupied by the pyramidal cells within the arc of the DG was considered a part of CA3 in the current study^44,45^. Areas of interest were selected using the interconnecting tool. Total optic density (o.d.,), expressed as digital light units per square millimeter (DLU/mm2), was measured over the area of interest using the OptiQuant software^10,46^, with the specific o.d. calculated using a threshold cutoff obtained from the white matter area of same immunostained section. We measured αSMA, BACE1, ZnT3 and COX immunolabeling in the MF field over the hilus and CA3 together. The pattern of immunolabeling of these markers around CA2 was disrupted with a great variability among the resected hippocampi, comparing to that in control hippocampal sections. It was difficulty to accurately define its borders to CA3 and CA1 in the surgical samples; therefore, we did not analyze the density of αSMA, BACE1, ZnT3 and COX IR in CA2 in the present study. Numeric densitometry was used to quantify NeuN and sortilin immunolabeled cells according to a method described earlier^47,48^. Briefly, the hippocampal and subicular regions were defined anatomically, including the hilus together with the CA3 (hilus/CA3), CA2, CA1, and the subicular subregions (Pro-S, Sub and Pre-S together). Using the coordinates (X and Y scales) of the image, a set of grids in 200×200 µm^2^ were randomly generated for each of area, including 10-15 grids for Hilus/CA3, CA1 and Sub, and 3-5 grids for the small-sized CA2 area. Neuronal somata were counted in each grid by experimenters who were blinded to the sample groups, with the neuronal density (number of cells per mm^2^) calculated for the measured hippocampal regions of each sample.

### Data processing, statistical analysis and figure preparation

Densitometric data were processed in Excel spreadsheets, including grouping, normalization and calculation of means and standard deviation. Considering the resected hippocampi were fixed following surgical removal while the control samples were fixed with postmortem delays, we normalized the optic density data to the overall means (i.e., including values from all areas from all samples) of the two sample groups, respectively, for each type of immunolabeling. The normalized data were imported into GraphPad spreadsheets (GraphPad Prism 10) and analyzed statistically using one-way analysis of variance (ANOVA) with *posthoc* tests, with P<0.05 set as the cutoff for significant difference. Figures were prepared with Photoshop 2024 by assembling representative micrographs and graphs from data analyses. Brightness and contrast adjustments were applied to microscopical images for proper visibility.

## Results

### Neuronal loss and granule cell dispersion in the resected hippocampi relative to control

Neuronal loss and granule cell dispersion are the key neuropathological criteria of MTLE; therefore we first assessed the pattern and extent of these changes in the resected hippocampi using NeuN immunolabeling for all neurons^4^ (Fig. 1), and sortilin immunolabeling for principal neurons^46,49^ (Fig. 2). Neuronal loss was found to have reached CA1 in all resected hippocampi with variability between samples and could be seen in the subicular subregions in severely sclerotized hippocampi (Fig. 1A-F; 2A-F). Labeled neuronal profiles were also reduced apparently in the hilus and CA3 area within the arc of the DG among most cases, and, to a lesser extent, in the CA3 segment outside DG. Neuronal profiles were better preserved in CA2 and the joining CA3 and CA1 areas in a given hippocampus (Fig. 1A-E; 2A-E). However, the remaining pyramidal neurons in this sector appeared to be misaligned, had increased intercellular space and shortened apical dendritic processes by closer examination, relative to their counterparts in control sections (Fig. 2D1, F1). Granule cell dispersion was evident in the resected hippocampi, with granule cells dislocated to the ML and formed bizarre-looking islands (Fig. 1B, B2; 2A). It should be noted that the NeuN-labeled neurons remained in CA1 were variable in size and appeared to be largely interneurons (Fig. 1C, C1). In comparison, the remaining sortilin-labeled neurons in this area had a relatively large soma with basal and apical dendritic processes, therefore appeared to represent the individually surviving pyramidal neurons (Fig. 2C, C1).

**Figure 1.**
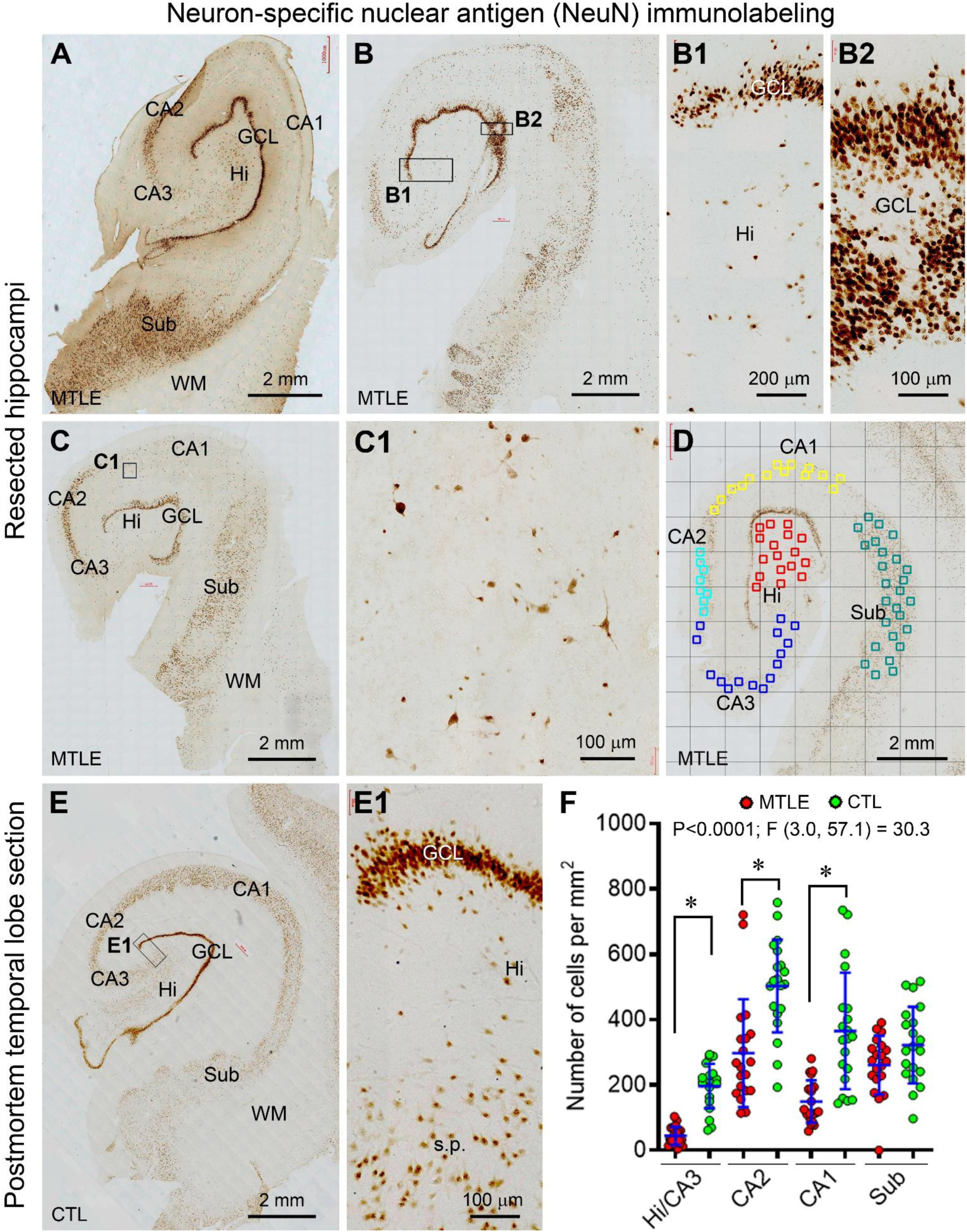
Assessment of neuronal loss in resected hippocampi from patients with mesial temporal lobe epilepsy (MTLE) in comparison with postmortem control with neuron-specific nuclear antigen (NeuN) immunohistochemistry. Shown are microscopical images from four resected hippocampi and a postmortem control (CTL), along with enlarged section areas as indicated. The labeled cellular profiles are greatly lost in CA1 in all resected samples (A, B, B1, C, C1, D) compared to CTL (E, E1), with neuronal loss also seen in the former in CA3 and subiculum (Sub) in varying extents (B, C). Granule cell dispersion is present in the resected hippocampi (A, B, B2, C), with ectopic granule cells forming bizarre-looking islands (B2). Pyramidal neurons in CA2 and the joining CA3 and CA1 areas are relatively preserved in the resected sections. Panel (D) shows the cell counting method used in this study, with the numerical densities of labeled neurons from the hilus (Hi) and CA3 combined at final data assembling. Statistically significant difference exists between the two sample groups for the numerical densities of labeled neurons in the hilus and CA3, CA2 and CA1 (F). Additional abbreviation: GCL: granule cell layer; s.p.: stratum pyramidale; WM: white matter. Scale bars are as indicated.

**Figure 2.**
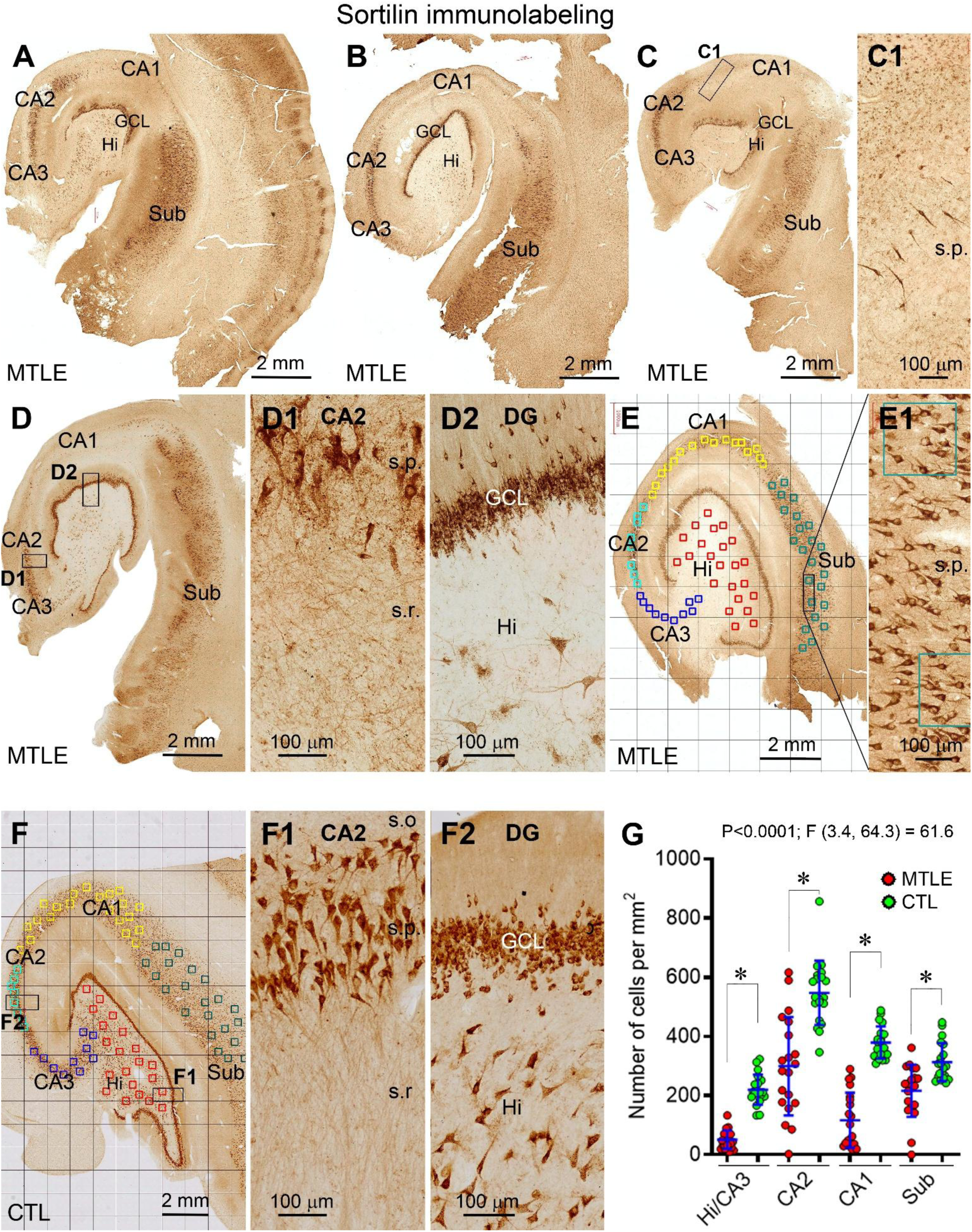
Assessment of neuronal loss in resected hippocampi relative to control with sortilin immunohistochemistry. Shown are microscopical images from five resected hippocampi from patients with mesial temporal lobe epilepsy (MTLE) (**A-E**) and a postmortem brain as control (CTL) (**F**). Loss of the labeled neurons is seen in the hilus (Hi), CA3 and CA1 (**A, B, C, D**) and also in the subiculum (Sub) with variability between samples (**B, C)**. Panel (**C1**) shows a few remaining pyramidal neurons in the striatum pyramidale (s.p.) of CA1. The pyramidal neurons in CA2 in the resected hippocampus are misaligned and separated from each other, and had fewer and shortened dendrites, in comparison with their counterpart in the control section (**D1, F1**). Dislocation of granule cells and loss of hilar mossy cells are clearly seen in the sclerotic (**D2**) as compared to the control (**F2**) hippocampus. Panels (**E, E1**) show cell counting in grided zones in the hippocampal subregions. Significant loss of sortilin-labeled neurons exists between the two sample groups in the hilus and CA3, CA2, CA1 and Sub (**G**). Additional abbreviation: GCL: granule cell layer; s.o.: stratum oriens; s.r.: stratum radiatum. Scale bars are as indicated.

Cell count was originally performed over the hilus, CA3, CA2, CA1, and the subicular subregions (Sub) together. However, the CA3 areas within and outside the hilar region were difficult to define in the resected hippocampi due to the cell loss. Also considering that neuronal loss was generally parallel in the hilar and CA3 areas among the resected samples, we combined the densities obtained from these subareas together in both MTLE and control groups at the final data assembly (Fig. 1F, 2G). The densities (mean±S.D.) of NeuN labeled profiles in the hilus/CA3 region were 44.5±27.6 cells/mm^2^ in the MTLE and 196.7±68.1 cells/mm^2^ in the control groups, respectively. The densities of NeuN labeled profiles were 297.7±165.1 vs. 502.9± 141.7 cells/mm^2^ in CA2, 149.4±64.7 vs. 365.3±178.1 cells/mm^2^ in CA1 and 274.6±67.8 vs. 322.2±116.9 cells/mm^2^ in Sub in the MTLE vs. control groups, respectively. There was an overall statistically significant difference of the means among the sample and region groups by one-way ANOVA [P<0.0001; F (DFn, DFd) = 29.3 (7, 151)]. Significant differences were reached between the two sample groups for the means in hilus/CA3, CA2, CA1, but not Sub, by *posthoc* Tukey’s multiple comparisons test (Fig. 1F). The extents of neuronal loss measured with sortilin immunolabeling in principal neurons showed a similar trend as with that of NeuN labeling. Thus, the numerical densities of sortilin-labeled neurons were 51.2±31.2 vs. 219.7± 51.0 cells/mm^2^ in hilus/CA3, 299.4±166.2 vs. 547.1±108.8 cells/mm^2^ in CA2, 116.9±93.3 vs.

379.3±54.5 cells/mm^2^ in CA1 and 227.4±74.2 vs. 312.8±64.1 cells/mm^2^ in Sub in the MTLE and control groups, respectively. There was also an overall difference in the means [P<0.0001; F (DFn, DFd) = 59.2 (7, 151)], with Tukey’s posthoc tests indicating statistically significant differences of the means in the hilus/CA3, CA2, CA1 and Sub regions between the two samples groups (Fig. 2G).

### Aberrant αSMA immunolabeling in the resected hippocampi relative to control

The neuropil-type, but not the vascular, αSMA immunoreactivity (IR) was altered in the resected hippocampi relative to postmortem controls (Fig. 3). We referred “αSMA IR” to this neuronal labeling exclusively in the following Result as well as the Discussion sections. The most prominent changes were the loss of αSMA IR in the hilus and CA3 regions, resulting in a spot-like remaining IR within and outside the arc of the DG (Fig. 3A1). On the contrary, the labeling was enhanced across the iML, with the somata of granule cells also labeled (Fig. 3A1, insert). The neuropil IR also appeared to be increased over the CA1 and subicular areas in the resected hippocampi relative to control (Fig. 3A-H), forming a band (pointed by arrows) extending from CA2, across the thinned CA1, and entering the subicular areas (Fig. 3A, A1, B-F). However, this band was much thinned and located deep to the stratum oriens (s.o.) in the heavily sclerotic hippocampi wherein the entire thickness of CA1 was dramatically narrowed (Fig. 3C, E).

**Figure 3.**
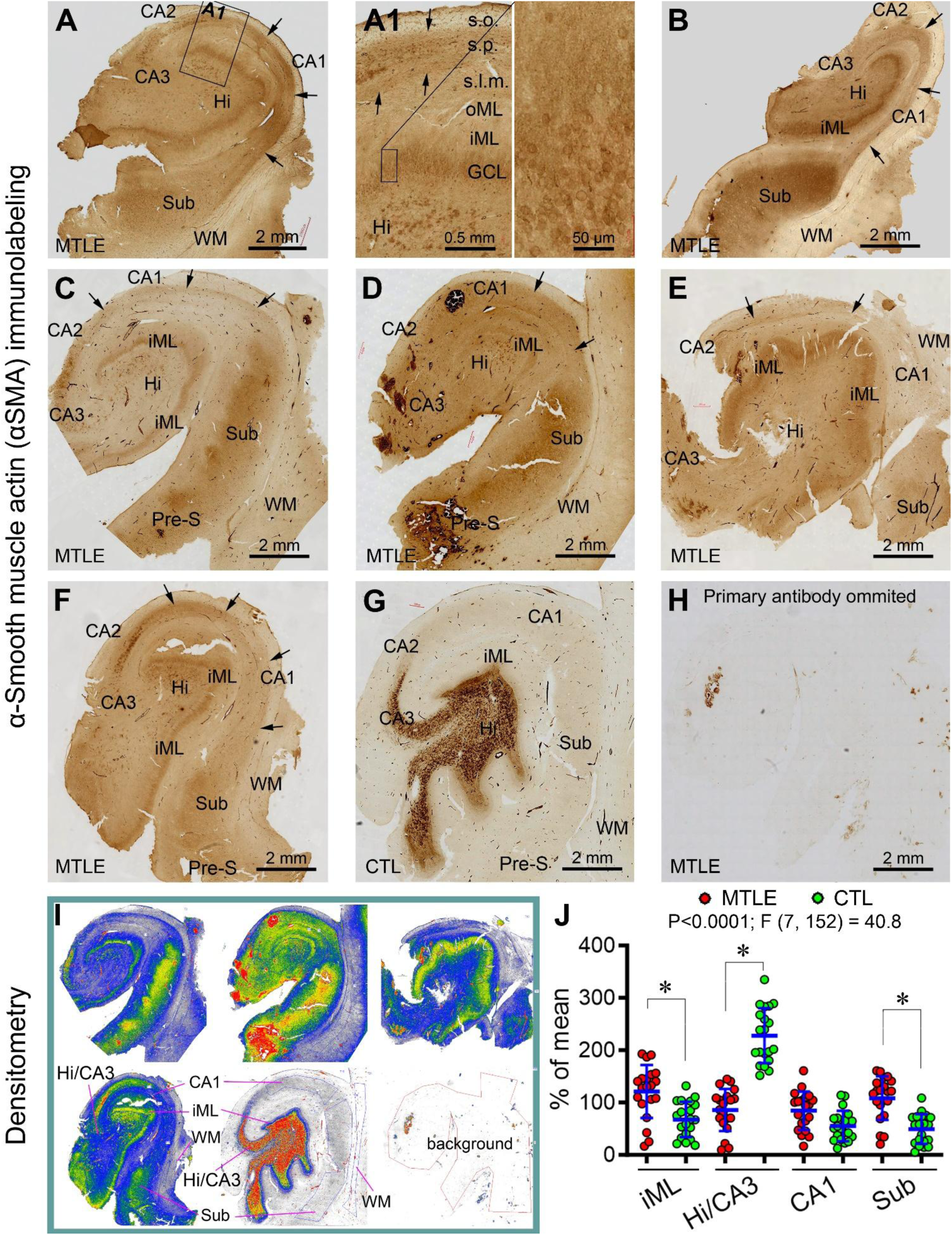
Aberrant ɑ-smooth muscle actin (αSMA) immunolabeling in resected hippocampi from patients with MTLE relative to postmortem control. Original micrographs show αSMA immunolabeled sections from 6 resected hippocampi (**A-F**), a postmortem control (**G**), and a parallelly processed section by excluding the primary antibody serving as assay control (**H)**. αSMA immunolabeling appears reduced in the hilus (Hi) and CA3, increased in the inner molecular layer (iML) and the subicular subregions (Sub) in the resected relative to control sections. The somata of the granule cells are visible in mix with the enhanced labeling of iML (**A1**). The labeling pattern around CA2 and its joining CA3 and CA1 areas appears disrupted in the resected samples (**A, A1, C, F**), as compared to the typical wadge-shape ending of the MF terminals in the control hippocampus (**G**). The labeling in CA1 appears as a band (pointed by arrows) in the resected hippocampi (**A, B, C, D, E, F**), which is flattened in the samples with the CA1 area greatly shrunken (**C, E**). Panel **(I)** illustrates densitometric method, with the areas of interest selected using the interconnecting drawing tool, avoiding blood clotting areas, torn/broken areas and large blood vessels. The optic density (o.d.) obtained from the white matter (WM) area is used as the cutoff for defining the specific o.d. in the areas of interest. The relative o.d. of αSMA labeling is significantly increased in the iML and Sub, reduced in the hilus/CA3 (**J**). Additional abbreviations: s.o.: stratum oriens; s.p.: stratum pyramidale; s.l.m.: stratum lacunosum-moleculare; Pre-S: presubiculum; WM: white matter. Scale bars are as indicated.

Optic densitometry for αSMA IR was carried out in hippocampal subregions using the white matter area in the same section as a background cutoff (Fig. 3I). Areas were circled using the free-hand interconnecting tool by excluding cracked areas, large blood vessels, and areas occupied by blood clots (showed non-specific labeling as seen in batch-processed sections excluding the αSMA antibody incubation, Fig. 3H). Given that the intensity of αSMA IR is similar in the hilus and CA3^41^ and it is difficult to define these subregions in the resected hippocampi, we measured the density over these areas together. In considering potential influence of tissue fixation delay, we calculated the relative densities of the regions by normalizing the raw data to the overall means (based on all regions from all samples) of the two groups, respectively. The relative densities (% of the overall mean) of αSMA IR in the iML were 121.2±50.6% (mean±S.D., same below) and 67.8±33.8% in the resected and control groups. The values were 86.0±40.3% vs. 227.7±52.5% in the hilus/CA3, 84.4±36.5% vs. 55.1±28.9% in the CA1 and 108.5±41.7% vs. 49.4±28.5% in the Sub regions, in the resected hippocampi vs. control groups, respectively. Statistically, there was an overall difference among the regional means [P<0.0001; F (DFn, DFd) = 40.8 (7, 152)]. Thus, the density of αSMA IR was increased significantly in the iML and Sub, but reduced in the hilus/CA3, areas in the resected group relative to control by Tukey’s multiple comparisons test, while no difference was reached for the density in CA1 between the two groups (Fig. 3J).

### Aberrant ZnT3 and BACE1 immunolabeling in the resected hippocampi relative to control

ZnT3 is a well-established marker for normal and sprouting hippocampal MF terminals^11,50^. BACE1 is the enzyme initiating the amyloidogenic pathway of the β-amyloid precursor protein (APP) proteolysis, which is enriched at the MF terminals and can mark MF sprouting in pilocarpine-induced rodent model of temporal lobe epilepsy^10,51^. We used these two markers to cross-validate the MF sprouting and to explore the extent thereof, if any, in the resected hippocampi, relative to that revealed by αSMA.

In comparison with control samples, ZnT3 IR was reduced in most subregions in the resected hippocampi, including the hilus/CA3 and CA1, and to a lesser extent, CA2 (Fig. 4A-F). The iML was widened in general but showed variability in its thickness as moving along the GCL, while its overall labeling intensity appeared to be similar or slightly enhanced relative to that seen at the same location in the control samples. The IR in CA1 and joining subicular subregions was greatly lost in some resected hippocampi, resulting in reduced intensity as low as that seen in the white matter (Fig. 4A, B, C). However, a thin band was visible near the s.o. even in these samples, similar to the αSMA band mentioned earlier. In other resected hippocampi, ZnT3 IR in the CA1 area appeared as a fused band, but it separated into two bands along the borders of the s.p. and merged into the subicular areas wherein these bands were no longer recognizable (Fig. 4D, E, E1). These two bands appeared to be anatomically consistent with infra- and supra-pyramidal bundles (IMF, SMF)^52^, which could be highlighted because of the loss of axonal terminals in the region. In postmortem control section, ZnT3 IR occurred across the entire CA2, CA1 and subicular subregions in an evenly distributed neuropil pattern (Fig. 4F). The relative densities of ZnT3 IR were obtained from the sections using the same densitometric approach as described for αSMA (Fig. 4G). The values in the iML did not reach significant difference between the resected (218.0±34.4%, mean±S.D.) and control (181.5±39.0%) groups. The densities of ZnT3 IR were reduced with statistically significant difference in the hilus/CA3 between the resected (136.3± 58.0%) vs. control (221.9±31.7%) groups, as well as in the CA1 area (115.8±57.3% vs. 174.1± 42.6%, same order). No statistical difference was reached for the values (203.0±23.6% vs. 177.0±36.1%) in the subicular areas (Fig. 4H).

**Figure 4.**
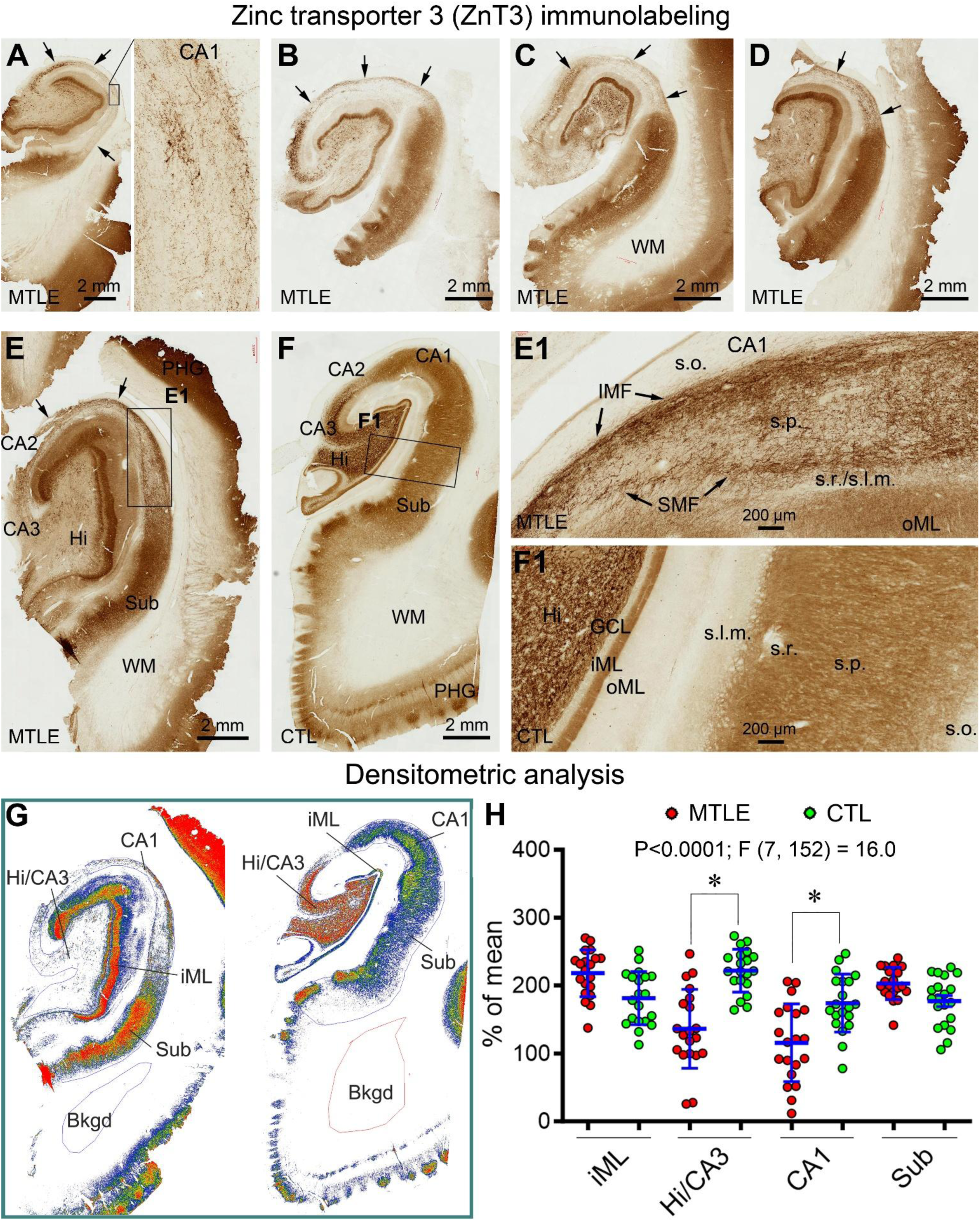
Aberrant Zinc transporter 3 (ZnT3) immunolabeling in resected hippocampi from patients with MTLE relative to postmortem control. Micrographs show altered labeling in five resected samples (**A, B, C, D, E, E1**) relative to a control (**F, F1**). ZnT3 immunolabeling is dramatically lost in the hilus/CA3 and CA1 areas in the resected hippocampi. The inner molecular layer (iML) is widened, with the labeling intensity somewhat enhanced, in the resected relative to control sections. In the resected samples, the labeling in CA2 is disrupted and appears to extend into CA1 and remains as across CA1 and further extending into the subicular regions; and as moving from CA1 towards the Sub, this band is separated into fibrous bundles more densely located around the borders of the stratum pyramidale (s.o.), which appear to be consistent anatomically with the so-called infra- and supra-pyramidal mossy fiber bundles (IMF, SMF, pointed by arrows) (**A** and enlarged area, **E, E1**). The immunolabeling in the preserved subicular areas and temporal cortex (in a subset of samples) in the resected hippocampi appears neuropil-like (**A-E**), as is the labeling across CA1, Sub and the temporal cortex in the postmortem control (**F, F1**). The densitometric method and results are demonstrated in (**G**) and (**H**), with statistically significant intergroup difference marked by asterisks (*). Abbreviations are as defined in Figure 3. Scale bars are as indicated.

An altered pattern of BACE1 IR was also seen in the resected hippocampi relative to control. Thus, BACE1 IR appeared to be increased in the iML, while there was an apparent loss of labeled MF terminals in the hilus and CA3 in the resected hippocampi (Fig. 5A-F). The remaining MF terminals in the hilus occurred in clusters along with lightly labeled mossy cells (Fig. 5E, E1). BACE1 IR appeared to be reduced in CA1 among most cases (Fig. 5A, C, D, E, F), but with a band remained deep to the s.o. in those with dramatic laminar thinning (Fig. 5C, E). The relative density of BACE1 IR was significantly increased in the iML (215.8±45.1% vs. 131.5±44.0%, resected vs. control, mean±S.D., same format below), reduced in the hilus/CA3 (135.3±60.6% vs. 222.4±51.7%), and reduced in CA1 (109.4±48.1% vs. 156.9±42.4%) in the resected relative to control groups. No significant intergroup difference was reached for the values (190.4±37.1% vs. 156.4±43.7%) in the subicular subregions (Fig. 5G, H).

**Figure 5.**
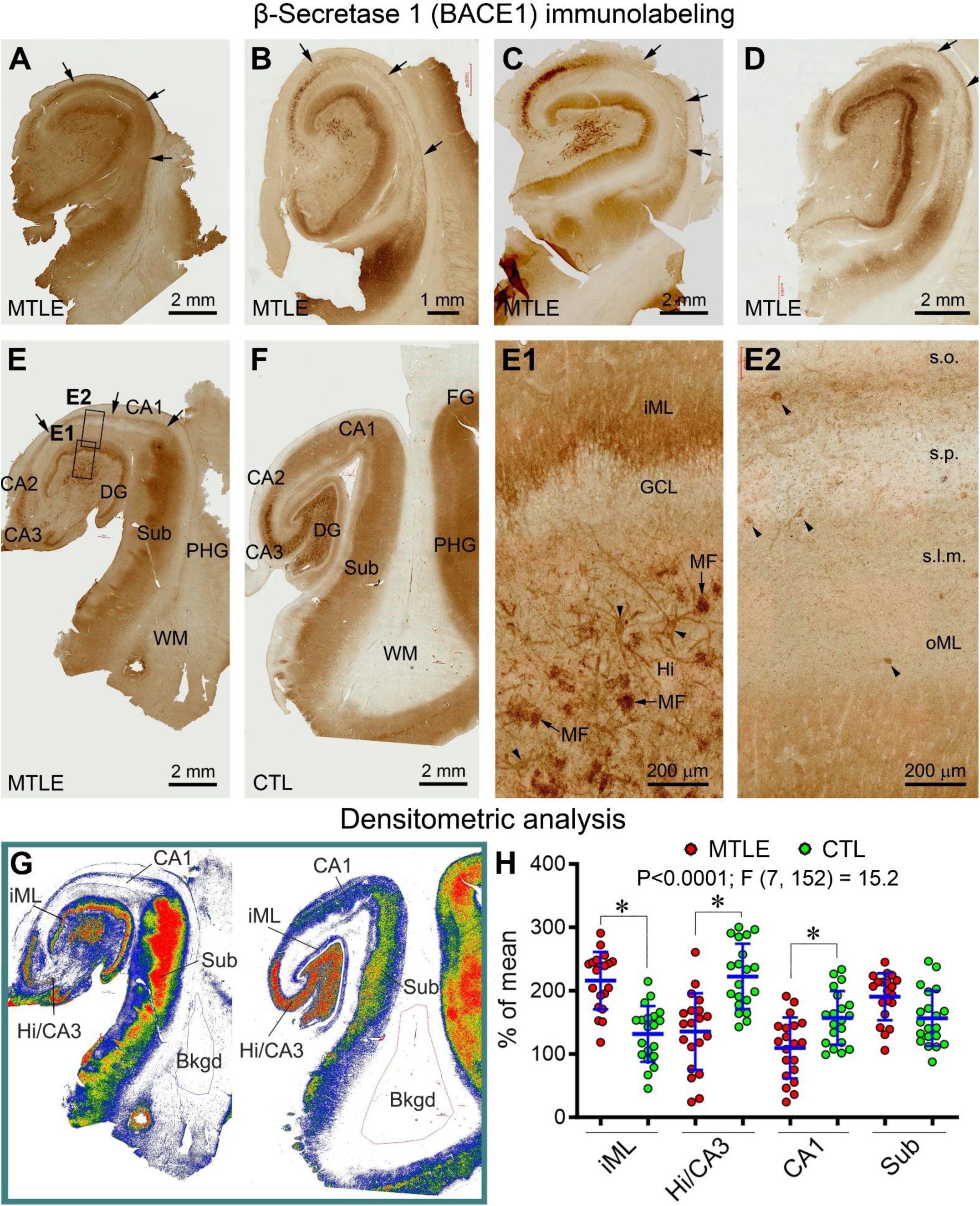
Aberrant β-secretase 1 (BACE1) immunolabeling in resected hippocampi from patients with MTLE relative to postmortem control. Panels (**A-F**) show low magnification images, with the framed areas enlarged as indicated (**E1, E2**). The mossy fiber (MF) labeling is reduced in the hilus and CA3, with the overall neuropil labeling in CA1 also reduced (with variability between cases) in the resected hippocampi (**A-E**) relative to control (**F**). The labeling in the subiculum appears to be increased in some (**B, E**), but reduced in other (**C, D**), resected hippocampi. However, a band-like labeling (arrows) remains in all resected hippocampi across CA1 and extending into the subiculum. Mossy cells in the hilus and some remaining interneurons (arrowheads) in CA1 show light BACE1 labeling (**E1, E2**). (**G**) shows the methodology for measuring specific optic densities in hippocampal subregions using the white matter area as a background (Bkgd) cutoff. (**H**) Graph and statistical results from one-way ANOVA analysis, with the * symbol indicating significantly intergroup difference. Scale bars are as indicated.

In the early phase of this study, we used sulphide fixation to prepare the resected hippocampi (a separate slice) along with fresh-froze postmortem human samples (n=4/group) for concurrent Timm stain and immunohistochemical analysis (Fig. S1 A-J). However, the background was increased in the immunolabeled sections with sulphide fixation (Fig. S1 B, C, B1, C1, F, G, I, J). In addition, while the Timm stain could reveal the MF changes in the dentate gyrus in the resected hippocampi relative to control, it could not visualize the neuropil alterations seen in the CA and subicular regions as described above in αSMA, ZnT3 and BACE1 preparations (Fig. S2). Therefore, this histological approach was no longer used in the following experiments.

The MF expression of αSMA in the hippocampus exhibits an evolutionary trend among mammals as such this labeling is not detectable in mice and rats, is visible in the CA3 area in guinea pigs and cats, but is distinctly present in CA3 and the dentate hilus in monkeys and humans^41^. In the present study we explored whether MF αSMA expression could be induced in experimental rodent model of temporal lobe epilepsy. C57BL/6 mice surviving 2 (n=4) months following pilocarpine-induced epileptic status along with vehicle-treated controls (n=4) were perfused with sulphide, rinsed with PBS and further fixed with 4% paraformaldehyde. Adjacent temporal lobe sections were prepared for Timm stain and immunohistochemistry. In Timm stain, MF sprouting into the iML occurred in the hippocampus of epileptic relative to control mice, with some darkly stained neuronal profiles also seen in CA1 and subiculum (Fig. S3 A, B). Neuropeptide Y (NPY) immunolabeling was dramatically enhanced in the dentate hilus and CA3 in the epileptic relative control hippocampal sections (Fig. S3 C, D). Further, BACE1 immunolabeling was increased in the iML, CA1 and subiculum in the epileptic hippocampus (Fig. S3 E, F). In contrast to the above markers, αSMA immunolabeling was only present in vascular profiles and did not visualize MFs in the hippocampi of control and pilocarpine-induced epileptic mice (Fig. S3 G, H).

### Aberrant COX immunolabeling in the resected hippocampi relative to control

Given that neuronal hyperactivity is the pathophysiological feature of temporal lobe epilepsy, we further explored potential regional alteration in mitochondrial metabolism in resected hippocampi relative to postmortem control using COX immunohistochemistry. Consistent with the histochemical staining pattern of this mitochondrial oxidative metabolism maker^53,54^, COX IR exhibited a patchy distribution pattern (blobs) in layers III-IV in the primary visual cortex of postmortem human brains (Fig. 6A, A1). Further, cortical interneurons exhibited heavy immunolabeling by close examination (Fig. 6A1 enlarged field), which was consistent with previously reported observations in COX histochemistry^55,56^. The regional and laminal distribution of COX IR was more differential in the resected hippocampi as compared with control (Fig. 6B-G). Thus, COX IR was heaviest in the iML and subicular areas in the resected hippocampi, which was related largely to an enhanced neuropil labeling (Fig. 6F, F1, F3). On the other hand, COR IR was reduced apparently in CA1, with the remaining labeling localized to interneurons and their dendritic processes (insert in Fig. 6E and Fig. 6F, F2). The relative densities of COX IR were significantly altered over the subregions in the resected hippocampi relative to control [P<0.0001, F (DFn, DFd) = 11.3 (9, 190)]. According to posthoc tests, the values were significantly increased in the MTLE relative to control groups in the iML (157.2±23.8% vs. 125.4±32.1%) and Sub (147.4±22.5% vs.116.6±25.3%), and reduced in CA1 (77.0±35.4%±119.7±22.5%), while no intergroup differences were reached for the outer molecular layer (oML) (115.7±33.9% vs. 116.7 ±26.2%) and the hilus/CA3 area (102.7±37.7% vs. 116.6±28.6%) (Fig. 6H).

**Figure 6.**
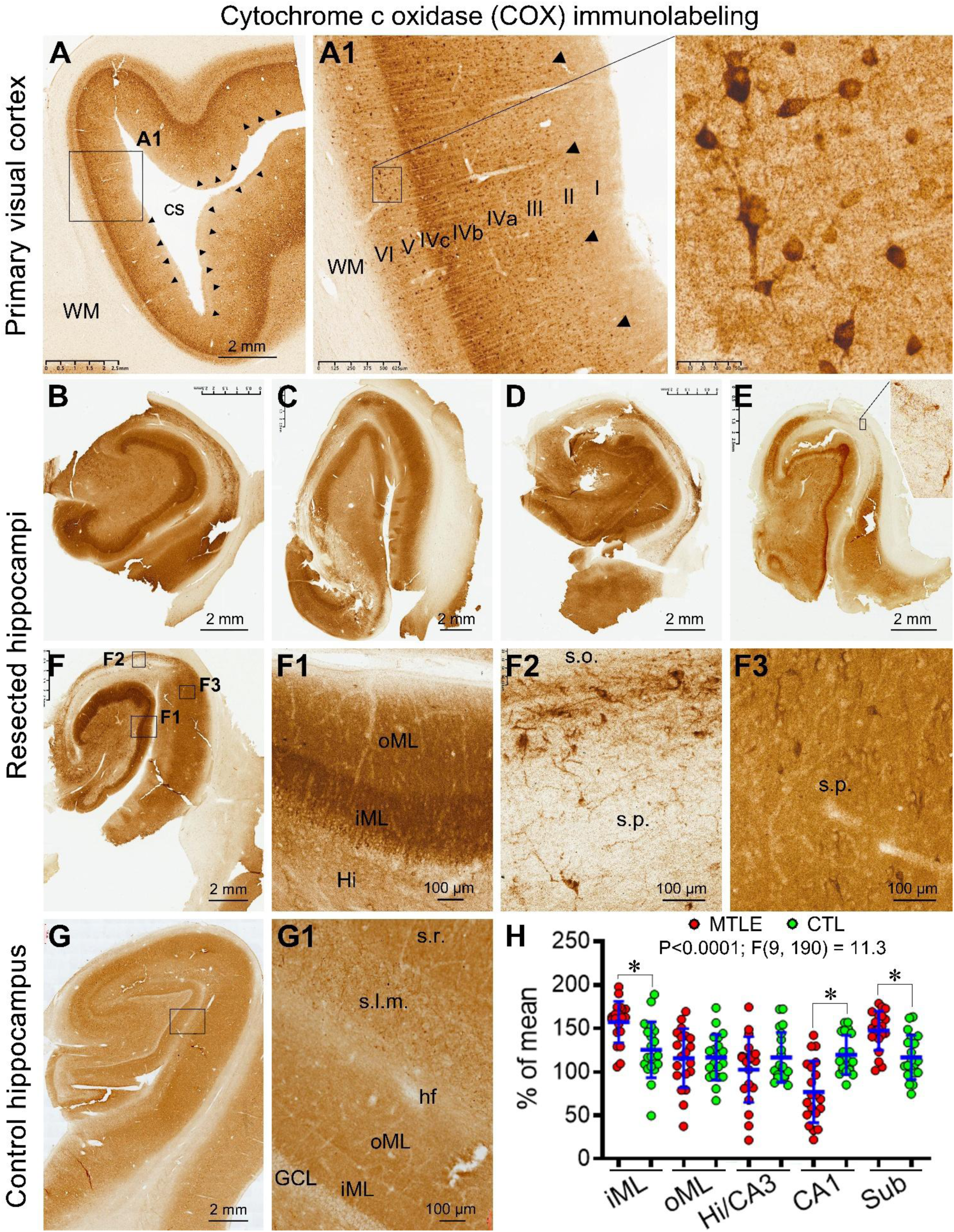
Characterization of cytochrome c oxidase (COX) immunolabeling in the primary visual cortex and its alteration in resected hippocampi from patients with MTLE relative to postmortem control (CTL). Panels **(A, A1)** show COX immunolabeling in the primary visual cortex from an adult human brain. The characteristic patchy pattern in the supragranular layers is displayed, with the columnar blobs indicated by arrowheads. The labeled cells appear to be largely interneurons, with the most heavily labeled ones being likely the basket cells (**A1** and enlarged view). Panels (**B-F**) and enlarged fields (**F1, F2, F3**) show altered labeling in the resected hippocampi as compared with control (**G, G1**). The distribution and intensity of COX labeling exhibit a great variability in the resected comparing to control hippocampal formation. The labeling appears lost over the CA1 area, and fairly intense in the inner molecular layer (iML) and subiculum (Sub) in the resected samples (**F, F1, F3**). The labeled cells remaining in CA1 appear to be interneurons according to their morphology (**F2**), with some pyramidal neurons labeled in the subiculum (**F3**). Quantitatively, the relative densities of COX labeling are significantly (*) decreased in CA1 and increased in the iML and subiculum between the resected and control sample groups (**H**). Abbreviations: cs: calcarine sulcus; I-VI: cortical layers; WM: white matter; oML: outer molecular layer; s.o.: stratum oriens; s.p.: stratum pyramidale; s.l.m.: stratum lacunosum-moleculare; hf: hippocampal fissure. Scale bars are as indicated.

## Discussion

The hippocampus appears to be the major epileptogenic center in drug-resistant MTLE, as such surgical resection of this structure is used as a last solution to control seizure recurrence^57–59^. Aberrant interictal electrographic activities can be detected in the hippocampus in patients with MTLE^60^, with dysfunctional communication existing between the hippocampus and extrahippocampal regions^61,62^. The cortical-hippocampal-cortical loop is a fundamental information processing system in the brain under normal conditions^63^, which appears to be disrupted at least on two major relays of the trisynaptic circuitry, i.e., the hilus/CA3 and CA1, in MTLE as a result of neuronal death. The preserved dentate granule cells along with their reorganized mossy fiber connections are commonly considered to generate and output neuronal hyperactivity^19,28,64–67^, while the subiculum may play a key role in the formation and propagation of epileptic activities^31,34,37,39,68^. The pathophysiology of MTLE remains puzzling in many aspects, with the anatomical/functional reorganization and interplay among the preserved neuronal populations, including those in the DG, CA2 and subiculum, potentially playing a critical role^20,69^.

The resected hippocampi examined in the current study had significant neuronal loss. Thus, the numerical densities of NeuN-labeled neurons were reduced significantly in the hilus/CA3 and CA1, and also in CA2, although did not reach significant difference in the Sub, in the resected hippocampi relative to control. Similarly, the densities of sortilin labeled neurons were significantly reduced in the resected samples in the hilus/CA3, CA1 and CA2, and also in Sub. The sortilin antibody marks largely the principal neurons^46^, which appeared more greatly lost in CA1 and Sub relative to the NeuN-labeled neurons in the resected hippocampi. These observations are in line with a severer loss of the principal neurons than interneurons in the hilus, CA1 and subicular regions^6,70–73^.

MF sprouting into the iML is consistent pathological feature in the hippocampus of MTLE patients and chronic epilepsy animal models as revealed by Timm stain and immunolabeling of multiple presynaptic synaptic markers^5–13^. Notably, studies have also shown that MF sprouting can extend into CA2^5,64,74^. In fact, in a recent study of 319 sclerotic hippocampi from patients, synaptoporin labeled axon terminals were observed in CA2, and even in CA1 in the specimens with relatively severe neuronal loss and granule cell dispersion^75^. Using retrovirus-based labeling technologies, MF sprouting into CA2 was observed in rats following pilocarpine-induced status epilepticus^76^ and in mice following kainate-induced chronic seizures^77^. This pattern was also observed in epileptic transgenic mice expressing enhanced green fluorescent protein^75^. Long range MF projection into the “regio superior” or CA1 appears to exist naturally among some mammalian species such as hedgehog and cats, with aberrant sprouting reported in cats with chronic epilepsy^52,78–80^.

In the present study we observed reduced αSMA, ZnT3 and BACE1 immunolabeling in the hilus/CA3 in the resected hippocampi relative to control, consistent with a loss of MF and likely other axon terminals in these areas. On the other hand, the iML was widened with strong αSMA, ZnT3 and BACE1 labeling in the resected hippocampi, consistent with MF sprouting into this lamina. αSMA, ZnT3 and BACE1 IR over the CA1 and subicular areas exhibited non-parallel alterations in the resected hippocampi. Specifically, αSMA IR was enhanced in CA1 and subiculum in the resected hippocampi relative to control. BACE1 and ZnT3 IR were reduced in CA1 in the resected hippocampi, which should be related to the loss of presynaptic axonal terminals associated with the neuronal loss in this region. However, a fibrous band was present in CA1 with αSMA, ZnT3 and BACE1 immunolabeling, which extended from CA2, across CA1 and entering the subicular areas. This band was flattened in the CA1 portion towards CA2 direction, and separated into fibrous bundles as moving towards the subicular areas. In the later sector, the tangentially running fibers were distributed more densely along the borders of the remaining pyramidal cell layer, which appeared to represent the so-called infrapyramidal and suprapyramidal bundles of the mossy fibers^52^. It should be noted that the above interpretations are based on the altered immunolabeling patterns of the MF-related markers. Apparently, we could not rule out that observed pattern changes are related to altered expression of the markers in local or distant cellular components. Nonetheless, there were few αSMA, ZnT3 and BACE1 immunolabeled somal and dendritic profiles in CA2, CA1 or subicular subregions in the resected hippocampi. In addition, no αSMA expression was observed in the sprouting MF in the mouse model of temporal lobe epilepsy induced by pilocarpine. It appears that ectopic or inducive neuronal aSMA expression may not, at least not commonly, occur the adult mammalian brain.

Neuronal firing is ATP-dependent^81^, therefore it is reasonable to expect that neuronal overexcitability in epileptogenic hippocampus would rely on high capacity of mitochondrial respiration. In this context, electron microscopic examination has found abundant presence of mitochondria in the reorganized MF terminals in the iML of human hippocampi^19^. Two previous studies (to the best of our knowledge) examined COX enzymatic activity with histochemistry in resected hippocampi from patients with MTLE with relatively small sample sizes (n=11, in reference^82^; and n=3, in reference^83^). Thus, COX activity was lost in the regions with cell loss, especially the CA1 area, while the enzyme activity was preserved^82^ or unaffected^83^ in the dentate molecular layer, CA2 and subiculum. Considering the impact of formalin fixation on histochemical enzyme reaction, immunohistochemistry was used in the present study to locate COX expression. The laminar and cellular pattern of COX IR in the primary visual cortex from postmortem human brains were consistent with established data^53–56^. With this cross-validation, we observed significantly elevated relative density of COX IR in the iML of DG and the subiculum, and reduced density in CA1.

## Conclusion

MF sprouting beyond the iML and pathological changes in the preserved hippocampal regions have gained increasing attention in study of MTLE in humans and animal models. The present study characterized the alteration of αSMA, ZnT3, BACE1 and COX immunolabeling in the hippocampi from MTLE patients with confirmed neuronal loss. These ML-related markers labeled MF sprouting in the iML. αSMA labeling was increased in the CA1 and Sub, while ZnT3, BACE1 labeling in CA1 were preserve in CA1 as a fibrous band extending into the Sub. Heavy COX immunolabeling occurred in the iML and subiculum. These findings suggest the existence of putative long range mossy fiber sprouting and regional high compacity of mitochondrial respiration in epileptogenic human hippocampus.

## Acknowledgements

We thank Jian Jiang, Jia-Qi Ai, Yan Wang, Xiao-Lu Cai and Sidiki Coulibaly for human brain banking and histopathological evaluation, and Xiao-Hua Tan, Ming-Li Zhang for help with light microscopic imaging.

## Availability of data and material

All data needed to evaluate the conclusions in the paper are present in the paper and/or the Supplementary Materials. The raw data will be made available by the authors by contacting Tian Tu and Xiao-Xin Yan, without undue reservation.

**Additional information:** Supplemental figures 1-3.

## Authors’ contributions

All authors read and approved the final version of the manuscript. Conceptualization and study design: XXY, ZYL; Methodology: TT, LW, CY, HSZ; Data acquisition and analysis: QLZ, ZPS, TT; Writing - original draft preparation: TT, XXY; Writing - review and editing: XXY; Funding acquisition: XXY, TT, AP, ET, ZYL; Resources: HYL, BST, ET, JW, ZQW, ZYL.

## Ethics approval

The surgical procedure and pathological examination of resected brain tissues were carried out with written consent from the patients and family members. All human brains were banked with written informed consent obtained prior to post-mortem donation from the donor and/or the next-of kin. The Ethics Committee for Research and Education at Xiangya School of Medicine approved the use of postmortem human materials.

## References

1. Téllez-Zenteno, J. F. & Hernández-Ronquillo L. A review of the epidemiology of temporal lobe epilepsy. Epilepsy Res Treat. 2012, 1–5(2012).

2. Barba, C. A. O. et al. Temporal lobe epilepsy surgery in children and adults: A multicenter study. Epilepsia 62, 128–142(2021).

3. Thom, M. Review: Hippocampal sclerosis in epilepsy: A neuropathology review. Neuropathol. Appl. Neurobiol. 40, 520–543(2014).

4. Blümcke, I. et al. International consensus classification of hippocampal sclerosis in temporal lobe epilepsy: A task force report from the ilae commission on diagnostic methods. Epilepsia 54, 1315–1329(2013).

5. Houser, C. R. et al. Altered patterns of dynorphin immunoreactivity suggest mossy fiber reorganization in human hippocampal epilepsy. J. Neurosci. 10, 267–282(1990).

6. Blümcke, I. et al. Temporal lobe epilepsy associated up-regulation of metabotropic glutamate receptors: Correlated changes in mglur1 mrna and protein expression in experimental animals and human patients. J. Neuropathol. Exp. Neurol. 59, 1–10(2000).

7. Proper, E. A. et al. Immunohistochemical characterization of mossy fibre sprouting in the hippocampus of patients with pharmaco-resistant temporal lobe epilepsy. Brain 123, 19–30(2000).

8. Furtinger, S. et al. Plasticity of y1 and y2 receptors and neuropeptide y fibers in patients with temporal lobe epilepsy. J. Neurosci. 21, 5804–5812(2001).

9. Van der Hel, W. S. et al. Hippocampal distribution of vesicular glutamate transporter 1 in patients with temporal lobe epilepsy. Epilepsia 50, 1717–1728(2009).

10. Yan, X. X. et al. Bace1 elevation is associated with aberrant limbic axonal sprouting in epileptic cd1 mice. Exp. Neurol. 235, 228–237(2012).

11. Crèvecœur, J. et al. Expression pattern of synaptic vesicle protein 2 (sv2) isoforms in patients with temporal lobe epilepsy and hippocampal sclerosis. Neuropathol. Appl. Neurobiol. 40, 191–204(2014).

12. Schmeiser, B., Zentner J., Prinz M., Brandt A. & Freiman T. M. Extent of mossy fiber sprouting in patients with mesiotemporal lobe epilepsy correlates with neuronal cell loss and granule cell dispersion. Epilepsy Res. 129, 51–58(2017).

13. Prada Jardim, A., et al. Characterising subtypes of hippocampal sclerosis and reorganization: Correlation with pre and postoperative memory deficit. Brain Pathol. 28, 143–154(2018).

14. Ribak, C. E. & Dashtipour K. Neuroplasticity in the damaged dentate gyrus of the epileptic brain. Prog. Brain Res. 136, 319–328(2002).

15. Dudek, F. E. & Sutula T. P. Epileptogenesis in the dentate gyrus: A critical perspective. Prog. Brain Res. 163, 755–773(2007).

16. Pallud, J. et al. Dentate gyrus and hilus transection blocks seizure propagation and granule cell dispersion in a mouse model for mesial temporal lobe epilepsy. Hippocampus 21, 334–343(2011).

17. Nadler, J. V. The recurrent mossy fiber pathway of the epileptic brain. Neurochem. Res. 28, 1649–1658(2003).

18. Zhang, W., Thamattoor A. K., LeRoy C. & Buckmaster P. S. Surviving mossy cells enlarge and receive more excitatory synaptic input in a mouse model of temporal lobe epilepsy. Hippocampus 25, 594–604(2015).

19. Zhang, N. & Houser C. R. Ultrastructural localization of dynorphin in the dentate gyrus in human temporal lobe epilepsy: A study of reorganized mossy fiber synapses. J. Comp. Neurol. 405, 472–490(1999).

20. Houser, C. R. Hippocampal sclerosis in temporal lobe epilepsy: New views and challenges. Jasper’s Basic Mechanisms of the Epilepsies 15–34(2024).

21. Okazaki, M. M., Molnár P. & Nadler J. V. Recurrent mossy fiber pathway in rat dentate gyrus: Synaptic currents evoked in presence and absence of seizure-induced growth. J. Neurophysiol. 81, 1645–1660(1999).

22. Gabriel, S. et al. Stimulus and potassium-induced epileptiform activity in the human dentate gyrus from patients with and without hippocampal sclerosis. J. Neurosci. 24, 10416–10430(2004).

23. Buckmaster, P. S. & Lew F. H. Rapamycin suppresses mossy fiber sprouting but not seizure frequency in a mouse model of temporal lobe epilepsy. J. Neurosci. 31, 2337–2347(2011).

24. Sloviter, R. S., Zappone C. A., Harvey B. D. & Frotscher M. Kainic acid-induced recurrent mossy fiber innervation of dentate gyrus inhibitory interneurons: Possible anatomical substrate of granule cell hyperinhibition in chronically epileptic rats. J. Comp. Neurol. 494, 944–960(2006).

25. Puhahn-Schmeiser, B., Leicht K., Gessler F. & Freiman T. M. Aberrant hippocampal mossy fibers in temporal lobe epilepsy target excitatory and inhibitory neurons. Epilepsia 62, 2539–2550(2021).

26. Spigelman, I. et al. Dentate granule cells form novel basal dendrites in a rat model of temporal lobe epilepsy. Neuroscience 86, 109–120(1998).

27. Zhou, Q. G. et al. Chemogenetic silencing of hippocampal neurons suppresses epileptic neural circuits. J. Clin. Invest. 129, 310–323(2019).

28. Sparks, F. T. et al. Hippocampal adult-born granule cells drive network activity in a mouse model of chronic temporal lobe epilepsy. Nat. Commun. 11, 6138(2020).

29. Lybrand, Z. R. et al. A critical period of neuronal activity results in aberrant neurogenesis rewiring hippocampal circuitry in a mouse model of epilepsy. Nat. Commun. 12, 1423(2021).

30. Hendricks, W. D., Westbrook G. L. & Schnell E. Early detonation by sprouted mossy fibers enables aberrant dentate network activity. Proc. Natl. Acad. Sci. U S A 116, 10994–10999(2019).

31. Cohen, I., Navarro V., Clemenceau S. P., Baulac M. & Miles R. On the origin of interictal activity in human temporal lobe epilepsy in vitro. Science 298, 1418–1421(2002).

32. Staba, R. J., Wilson C. L., Bragin A., Fried I. & Engel J. Quantitative analysis of high-frequency oscillations (80–500 hz) recorded in human epileptic hippocampus and entorhinal cortex. J. Neurophysiol. 88, 1743–1752(2002).

33. Vreugdenhil, M., Hoogland G., Van Veelen C. W. M. & Wadman W. J. Persistent sodium current in subicular neurons isolated from patients with temporal lobe epilepsy. Eur. J. Neurosci. 19, 2769–2778(2004).

34. Fabó, D. et al. Properties of in vivo interictal spike generation in the human subiculum. Brain 131, 485–499(2008).

35. Wozny, C., Knopp A., Lehmann T. N., Heinemann U. & Behr J. The subiculum: A potential site of ictogenesis in human temporal lobe epilepsy. Epilepsia 46, 17–21(2005).

36. Huberfeld, G. et al. Glutamatergic pre-ictal discharges emerge at the transition to seizure in human epilepsy. Nat. Neurosci. 14, 627–634(2011).

37. Kitaura, H. et al. Pathophysiological characteristics associated with epileptogenesis in human hippocampal sclerosis. EBioMedicine 29, 38–46(2018).

38. Xu, C. et al. Subicular pyramidal neurons gate drug resistance in temporal lobe epilepsy. Ann. Neurol. 86, 626–640(2019).

39. Lévesque, M. & Avoli M. The subiculum and its role in focal epileptic disorders. Rev. Neurosci. 32, 249–273(2020).

40. Fei, F., Wang X., Wang Y. & Chen Z. Dissecting the role of subiculum in epilepsy: Research update and translational potential. Prog. Neurobiol. 201, 102029(2021).

41. Tu, T. et al. Mossy fiber expression of αsma in human hippocampus and its relevance to brain evolution and neuronal development. Sci. Rep. 15, 15834(2025).

42. Yan, X. X., Ma C., Bao A. M., Wang X. M. & Gai W. P. Brain banking as a cornerstone of neuroscience in china. Lancet Neurol. 14, 136(2015).

43. Qiu, W. et al. Standardized operational protocol for human brain banking in china. Neurosci. Bull. 35, 270–276(2019).

44. Amaral, D. G. & R I. Hippocampal formation. The Human Nervous System. *New York*: *Academic Press* 711–755(1990).

45. Amaral, D. G., Fau S. H. & Lavenex P. The dentate gyrus: Fundamental neuroanatomical organization (dentate gyrus for dummies). Prog. Brain Res. 163, 3–22(2007).

46. Xu, S. Y. et al. Regional and cellular mapping of sortilin immunoreactivity in adult human brain. Front. Neuroanat. 13, 31(2019).

47. Ai, J. Q. et al. Doublecortin-expressing neurons in chinese tree shrew forebrain exhibit mixed rodent and primate-like topographic characteristics. Front. Neuroanat. 15, 727883(2021).

48. Li, Y. N. et al. Doublecortin-expressing neurons in human cerebral cortex layer ii and amygdala from infancy to 100 years old. Mol Neurobiol. 60, 3464–3485(2023).

49. Hu, X. et al. Sortilin fragments deposit at senile plaques in human cerebrum. Front. Neuroanat. 11, 2017).

50. Wenzel, H. J., Cole T. B., Born D. E., Schwartzkroin P. A. & Palmiter R. D. Ultrastructural localization of zinc transporter-3 (znt-3) to synaptic vesicle membranes within mossy fiber boutons in the hippocampus of mouse and monkey. Proc. Natl. Acad. Sci. U S A 94, 12676–12681(1997).

51. Zhang, S., Khanna S. & Tang F. R. Patterns of hippocampal neuronal loss and axon reorganization of the dentate gyrus in the mouse pilocarpine model of temporal lobe epilepsy. J. Neurosci. Res. 87, 1135–1149(2009).

52. Gaarskjaer, F. The organization and development of the hippocampal mossy fiber system. Brain Res. 396, 335–357(1986).

53. Horton, J. C. & Hubel D. H. Regular patchy distribution of cytochrome oxidase staining in primary visual cortex of macaque monkey. Nature 292, 762–764(1981).

54. Wong-Riley, M. T. T. et al. Cytochrome oxidase in the human visual cortex: Distribution in the developing and the adult brain. Vis. Neurosci. 10, 41–58(1993).

55. Luo, X. G., Hevner R. F. & Wong-Riley M. T. T. Double labeling of cytochrome oxidase and γ-aminobutyric acid in central nervous system neurons of adult cats. J. Neurosci. Methods 30, 189–195(1989).

56. Nie, F. & Wong-Riley M. T. T. Double labeling of gaba and cytochrome oxidase in the macaque visual cortex: Quantitative em analysis. J. Comp. Neurol. 356, 115–131(1995).

57. Huberfeld, G., Blauwblomme T. & Miles R. Hippocampus and epilepsy: Findings from human tissues. Rev. Neurol. (Paris) 171, 236–251(2015).

58. Shi, W. et al. Spike ripples localize the epileptogenic zone best: An international intracranial study. Brain 147, 2496–2506(2024).

59. Dong, H., Shi J., Wei P., Shan Y. & Zhao G. Comparative efficacy of surgical strategies for drug-resistant epilepsy: A systematic review and meta-analysis. World Neurosurg. 195, 123729(2025).

60. Pail, M. et al. High frequency oscillations in epileptic and non-epileptic human hippocampus during a cognitive task. Sci. Rep. 10, 18147(2020).

61. Mendes, R. A. V. et al. Hijacking of hippocampal-cortical oscillatory coupling during sleep in temporal lobe epilepsy. Epilepsy Behav. 121, 106608(2021).

62. Hu, J. et al. Interictal suppression in patients with mesial temporal lobe epilepsy: A simultaneous pet/fmri study. Neuroimage 314, 121207(2025).

63. Rothschild, G., Eban E. & Frank L. M. A cortical–hippocampal–cortical loop of information processing during memory consolidation. Nat. Neurosci. 20, 251–259(2017).

64. Wittner, L. et al. The epileptic human hippocampal cornu ammonis 2 region generates spontaneous interictal-like activity in vitro. Brain 132, 3032–3046(2009).

65. Krook-Magnuson, E., et al. In vivo evaluation of the dentate gate theory in epilepsy. J. Physiol. 593, 2379–2388(2015).

66. Dengler, C. G., Yue C., Takano H. & Coulter D. A. Massively augmented hippocampal dentate granule cell activation accompanies epilepsy development. Sci. Rep. 7, 42090(2017).

67. Kahn, J. B., Port R. G., Yue C., Takano H. & Coulter D. A. Circuit-based interventions in the dentate gyrus rescue epilepsy-associated cognitive dysfunction. Brain 142, 2705–2721(2019).

68. de Guzman, P. et al. Subiculum network excitability is increased in a rodent model of temporal lobe epilepsy. Hippocampus 16, 843–860(2006).

69. Whitebirch, A. C. et al. Enhanced excitability of the hippocampal ca2 region and its contribution to seizure activity in a mouse model of temporal lobe epilepsy. Neuron 110, 3121–3138.e8(2022).

70. de Lanerolle, N. C., Kim J. H., Robbins R. J. & Spencer D. D. Hippocampal interneuron loss and plasticity in human temporal lobe epilepsy. Brain Res. 495, 387–395(1989).

71. Mathern, G. W., Babb Tl Fau - Vickrey B. G., Vickrey Bg Fau - Melendez M., Melendez M Fau - Pretorius J. K. & Pretorius J. K. The clinical-pathogenic mechanisms of hippocampal neuron loss and surgical outcomes in temporal lobe epilepsy. Brain : a journal of neurology 118, 105–118(1995).

72. Sperk, G., Hamilton T Fau - Colmers W. F. & Colmers W. F. Neuropeptide y in the dentate gyrus. Prog. Brain Res. 163, 285–297(2007).

73. Tóth, K. & Maglóczky Z. The vulnerability of calretinin-containing hippocampal interneurons to temporal lobe epilepsy. Front. Neuroanat. 8, 100(2014).

74. Cook, T. M. & Crutcher K. A. Lesion-induced ca1 mossy fibers in the rat represent a neoinnervation. Exp. Brain Res. 70, 433–436(1988).

75. Freiman, T. M. et al. Mossy fiber sprouting into the hippocampal regionca2in patients with temporal lobe epilepsy. Hippocampus 31, 580–592(2021).

76. Althaus, A. L., Zhang H. & Parent J. M. Axonal plasticity of age-defined dentate granule cells in a rat model of mesial temporal lobe epilepsy. Neurobiol. Dis. 86, 187–196(2016).

77. Häussler, U., Rinas K., Kilias A., Egert U. & Haas C. A. Mossy fiber sprouting and pyramidal cell dispersion in the hippocampal ca2 region in a mouse model of temporal lobe epilepsy. Hippocampus 26, 577–588(2016).

78. Gaarskjaer, F. B., Danscher G. & West M. J. Hippocampal mossy fibers in the regio superior of the european hedgehog. Brain Res. 237, 79–90(1982).

79. Laurberg, S. & Zimmer J. Aberrant hippocampal mossy fibers in cats. Brain Res. 188, 555–559(1980).

80. Blaabjerg, M. & Zimmer J. The dentate mossy fibers: Structural organization, development and plasticity. Prog. Brain Res. 163, 85–107(2007).

81. Li, S. & Sheng Z. H. Energy matters: Presynaptic metabolism and the maintenance of synaptic transmission. Nat. Rev. Neurosci. 23, 4–22(2021).

82. Brines, M. L. et al. Regional distributions of hippocampal na+, k+-atpase, cytochrome oxidase, and total protein in temporal lobe epilepsy. Epilepsia 36, 371–383(2005).

83. Opačić M. et al. Regional distribution of cytochrome c oxidase activity and copper in sclerotic hippocampi of epilepsy patients. Brain Behav. 11, e01986(2020).

