## Supplemental Figures 1-3 for "Putative long-range mossy fiber sprouting and regional hypermetabolic capacity in the hippocampus of patients with mesial temporal lobe epilepsy"

### Putative long-range mossy fiber sprouting and regional hypermetabolic capacity in the hippocampus of patients with mesial temporal lobe epilepsy

Tian Tu[1,2], Lily Wan[2], Qi-Lei Zhang[2], Chen Yang[2], Hong-Shu Zhou[3], Zhong-Ping Sun[2], Hong-Yu Long[1], Bei-Sha Tang[1], Aihua Pan[2], Ewen Tu[4], Jian Wang[5], Zhi-Quan Yang[3], Zhen-Yan Li[3*] and Xiao-Xin Yan[2*]

[1]Department of Neurology, Xiangya Hospital, Central South University, Changsha, Hunan 410008, China
[2]Department of Anatomy and Neurobiology, Xiangya School of Basic Medical Sciences, Central South University, Changsha, Hunan 410013, China
[3]Department of Neurosurgery, Xiangya Hospital, Central South University, Changsha, Hunan 410008, China
[4]Department of Neurology, The Second People's Hospital of Hunan Province, Changsha, Hunan 410007, China
[5]The Reproductive and Stem Cell Engineering Institute, Xiangya School of Basic Medical Sciences, Central South University, Changsha, Hunan 410013, China

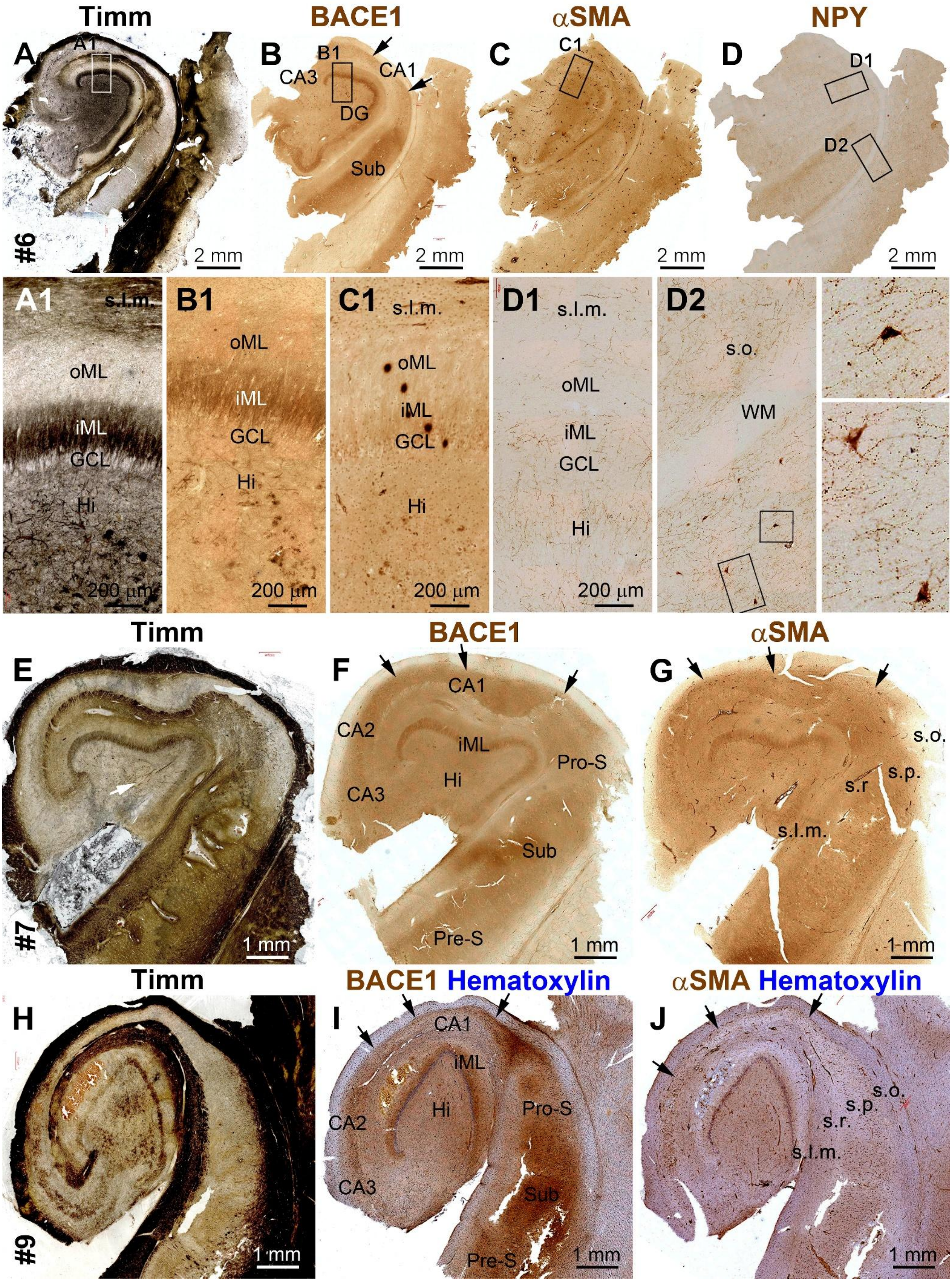
Timm
BACE1
αSMA
NPY
A
A1
#6
2 mm
B
B1
CA3
CA1
DG
Sub
C
C1
D
D1
D2
A1
s.l.m.
oML
iML
GCL
Hi
200 μm
B1
C1
D1
D2
s.o.
WM
E
F
G
CA1
CA2
CA3
iML
Hi
Pro-S
Sub
Pre-S
s.o.
s.p.
s.r
s.l.m.
#7
1 mm
Timm
BACE1 Hematoxylin
αSMA Hematoxylin
H
I
J
#9

**Supplemental Figure 1. Comparative assessment of Timm stain and immunolabeling of β-secretase 1 (BACE1), αSMA and neuropeptide Y (NPY) in sections of resected human hippocampal slices immerse-fixed with sulphide**

Low magnification images show the pattern of labeling in adjacent sections from three samples (A-D, E-G and H-J), with the framed areas enlarged as indicated **(A1, B1, C1, D1, D2)**. Timm stain labels mossy fiber sporting into the inner molecular layer (iML) and mossy fiber loss in the hilus in all resected hippocampi from patients with mesial temporal lobe epilepsy (**A, A1, E, H**). However, the staining is pale in the cellular layers of the hippocampal formation including the CA1 and subicular areas that exhibited significant alterations of aSMA, BACE1 and ZnT3 labeling using formalin without sulphide fixation as shown in the main figures. In these sulphide fixed tissues, there is a high background reactivity in BACE1 and aSMA immunolabeled sections, although the iML is strongly labeled (**B1, C1, F, G, I, J**). NPY immunolabeling is reportedly an excellent method for mossy fiber pathology in epileptic animal models. However, it did not well visualize the MF sprouting in the resected human hippocampi with sulphide (**D, D1, D2**) or formalin fixation (not shown) as assessed in the present study, although the antibody can clearly label the interneurons and their processes in the human brain sections (**D2**). Therefore, this antibody labeling method was not used in the following formal experiments with the human samples after the initial assessment as presented here. Abbreviations for neuroanatomical structures: CA1-3: subregions of the Ammon’s horn; DG: dentate gyrus; Sub: subiculum; Pro-S: prosubiculum; Pre-S: presubiculum; Hi: hilus; WM: white matter; s.o.: stratum oriens; s.p.: stratum pyramidale; s.l.m.: stratum lacunosum-moleculare; s.r.: stratum radiatum; oML: outer molecular layer; GCL: granule cell layer; Scale bars are as indicated.

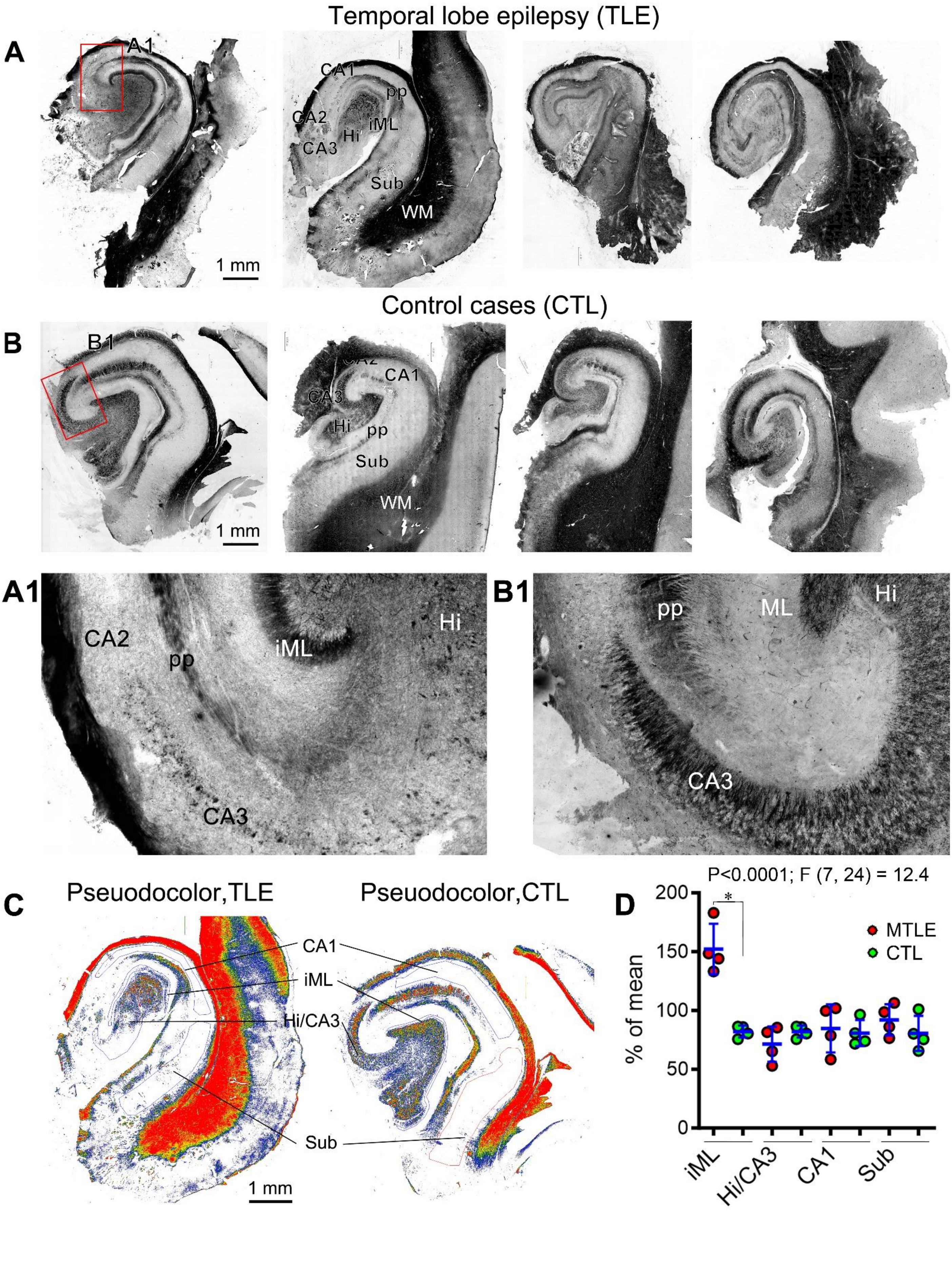
Temporal lobe epilepsy (TLE)
A
A1
CA1
pp
CA2
iML
Hi
CA3
Sub
WM
1 mm
Control cases (CTL)
B
B1
CA2
CA1
CA3
Hi
pp
Sub
WM
1 mm
A1
CA2
pp
iML
Hi
CA3
B1
pp
ML
Hi
CA3
C
Pseudodocolor,TLE
Pseuodocolor,CTL
CA1
iML
Hi/CA3
Sub
1 mm
P<0.0001; F (7, 24) = 12.4
D
MTLE
CTL
% of mean
0
50
100
150
200
*
iML
Hi/CA3
CA1
Sub

**Supplemental Figure 2. Timm stain and quantification of regional density in resected and control human hippocampi immerse-fixed with sulphide followed by formalin.** Low magnification images of Timm stained sections from four resected hippocampi (**A**) and the temporal lobe slices fresh-prepared from postmortem human brains (**B**). The framed areas in (**A, B**) are enlarged as (**A1**) and (**B1**). Mossy fiber sprouting into the inner molecular layer (iML) are present in the resected hippocampi as compared to controls (A1, B1). Dark zinc staining is present in the white matter of the temporal cortex and the major fiber pathways of the hippocampal formation including the alveus (Alv) and the perforant path (pp). The MF terminals in the hilus and CA3 are largely lost in the sclerotic hippocampi as compared with control (A1, B1). **(C）** Densitometric sampling in hippocampal subregions as indicated. (**D**) Graph and statistical results of densitometric analysis of Timm stain over different subregions, with the regional densities normalized to the mean of all regions from two sample groups, respectively. Statistically significant difference only existed for the density in the iML between the two groups. PP: Perforant path: Other abbreviations of anatomical structures are as defined in Supplemental Figure 1. Scale bars are as indicated.

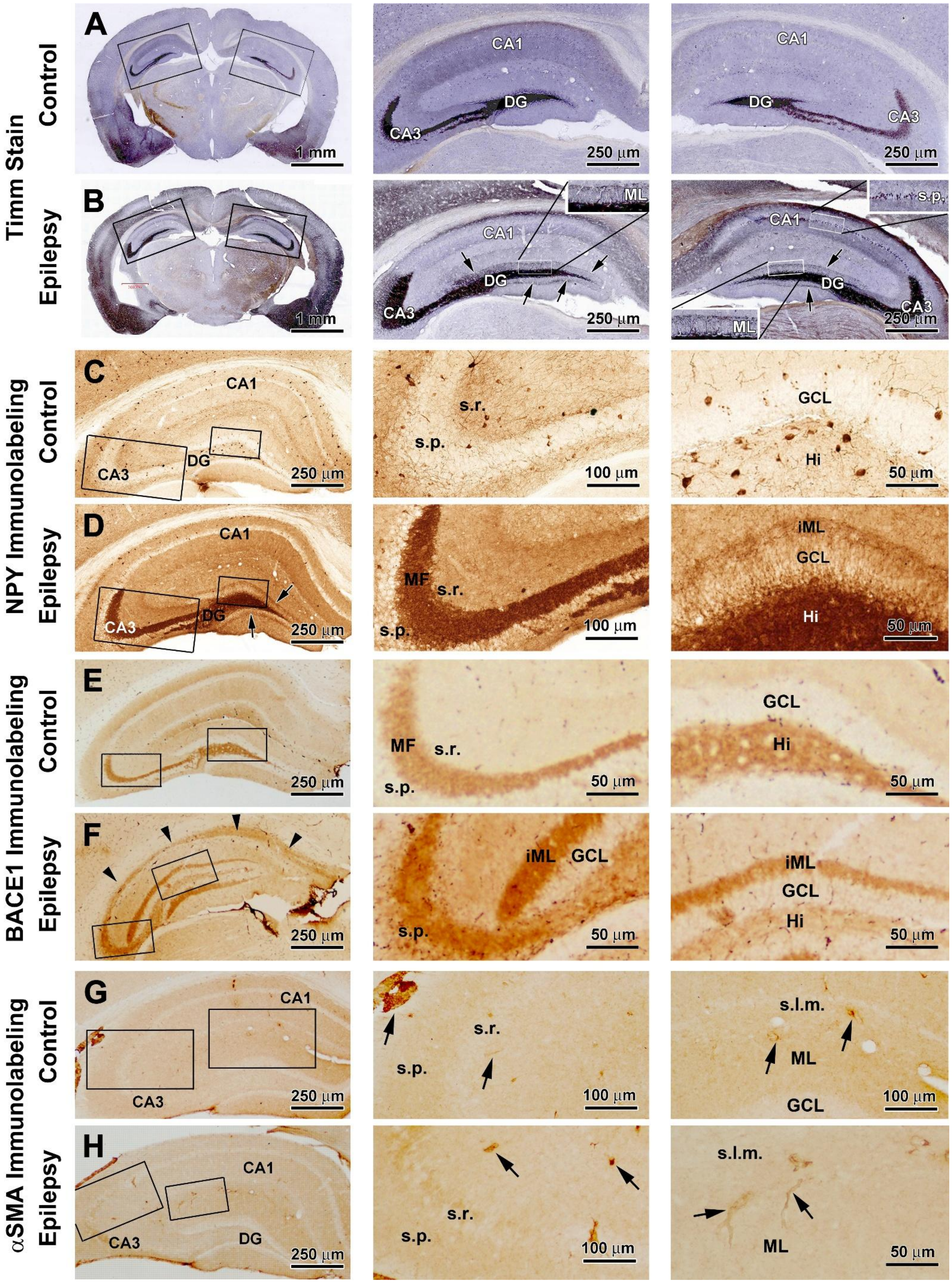

Timm Stain
Control
Epilepsy
NPY Immunolabeling
Control
Epilepsy
BACE1 Immunolabeling
Control
Epilepsy
αSMA Immunolabeling
Control
Epilepsy
A
CA1
DG
CA3
1 mm
250 µm
B
ML
s.p.
CA1
DG
CA3
1 mm
250 µm
C
CA1
DG
CA3
s.r.
s.p.
GCL
Hi
250 µm
100 µm
50 µm
D
CA1
DG
CA3
MF
s.r.
s.p.
iML
GCL
Hi
250 µm
100 µm
50 µm
E
MF
s.r.
s.p.
GCL
Hi
250 µm
50 µm
F
iML
GCL
s.p.
Hi
250 µm
50 µm
G
CA1
CA3
s.r.
s.p.
s.l.m.
ML
GCL
250 µm
100 µm
H
CA1
CA3
DG
s.r.
s.p.
s.l.m.
ML
250 µm
100 µm
50 µm

**Supplemental Figure 3. Representative micrographs showing a lack of αSMA expression in the hippocampal mossy fibers in control as well as epileptic mice induced by pilocarpine.** Terminal histological studies were performed 2 months after intraperitoneal injection of pilocarpine inducing acute seizures and chronic epileptic status, with the control received vehicle treatment. Staining methods and animal groups are as indicated. In Timm stain preparation (**A, B**), MF sprouting into the inner molecular layer (iML) occurred in both sides of the hippocampus in the epileptic relative to control samples, with the overall neuropil-like labeling also somewhat enhanced in the amygdala, basal temporal cortex, CA1 and subiculum, in the former. Some darkly stained neurons are visible at high magnification in the pyramidal cell layer of CA1 and subiculum in the epileptic sections (**B**, enlarged views). Neuropeptide Y (NPY) immunolabeling is only present in the interneurons and their processes in the control hippocampus (**C**). In the epileptic hippocampus, the mossy fibers in the dentate hilus and CA3 show heavy *de nova* NPY immunoreactivity, while light labeling also occurred in the iML **(D**). BACE1 immunolabeling reveals the normal MF distribution in the hilus and CA1 in the control section, with a light and even neuropil-like reactivity in the hippocampal layers and cortical grey matter (**E** and enlarged panels). In the epileptic hippocampus, iML is clearly labeled in the dentate gyrus. There is also a moderately enhanced BACE1 immunolabeling appearing as a band (pointed by arrows) over the stratum pyramidale extending from CA2 to CA1, and further to the subiculum (**F** and enlarged views). The αSMA immunolabeling is solely related to vascular profiles in both the control and epileptic hippocampal sections. Thus, no inducible aSMA expression occurs in sprouting MFs in rodent model of temporal lobe epilepsy. Abbreviations of anatomical structures are as defined in Supplemental Figure 1. Scale bars are as indicated.
